# Marine Fe-oxidizing *Zetaproteobacteria*: Historical, ecological, and genomic perspectives

**DOI:** 10.1101/416842

**Authors:** Sean M. McAllister, Ryan M. Moore, Amy Gartman, George W. Luther, David Emerson, Clara S. Chan

**Author notes:** Address correspondence to Clara S. Chan. Present address: Amy Gartman, USGS Pacific Coastal and Marine Science Center, Santa Cruz, California, USA.

## Abstract

The *Zetaproteobacteria* are a class of bacteria typically associated with marine Fe oxidizing environments. First discovered in the hydrothermal vents at Loihi Seamount, Hawaii, they have become model organisms for marine microbial Fe oxidation. In addition to deep sea and shallow hydrothermal vents, *Zetaproteobacteria* are found in coastal sediments, other marine subsurface environments, steel corrosion biofilms, as well as saline terrestrial aquifers and springs. Isolates from a range of environments all grow by Fe oxidation. Their success lies partly in their microaerophily, which enables them to compete with abiotic Fe oxidation at the low O_2_ concentrations common to Fe(II)-rich oxic/anoxic transition zones. Also, *Zetaproteobacteria* make a variety of biomineral morphologies as a repository for Fe(III) waste, and as attachment structures. To determine the known diversity of the *Zetaproteobacteria*, we have used 16S rRNA gene sequences to define 59 operational taxonomic units (OTUs), at 97% similarity. While some *Zetaproteobacteria* taxa appear to be cosmopolitan, various habitats enrich for different sets of *Zetaproteobacteria*. OTU networks show that certain *Zetaproteobacteria* co-exist, sharing compatible niches. These niches may correspond with adaptations to O_2_, H_2_, and nitrate availability, based on genomic analyses. Also, a putative Fe oxidation gene has been found in diverse *Zetaproteobacteria* taxa, suggesting that the *Zetaproteobacteria* evolved as specialists in Fe oxidation. In all, culture, genomic, and environmental studies suggest that *Zetaproteobacteria* are widespread, and therefore have a broad influence on marine and saline terrestrial Fe cycling.

## Introduction

Fe in marine environments is a study in contrasts. It is often a limiting nutrient in the open ocean, while the basaltic ocean crust and many sediments are replete in Fe. This stark difference is due to the redox chemistry of Fe, which is present as Fe(II) in basalt and anoxic groundwater, but rapidly oxidizes to Fe(III) in oxic ocean water, precipitating as Fe(III) minerals. This oxidation was assumed to be dominated by rapid abiotic oxidation at circumneutral pH, but the discovery of the marine Fe-oxidizing *Zetaproteobacteria* gave proof that the process can be driven by microbes. First proposed as a class in 2007 (1), *Zetaproteobacteria* have since been widely observed in deep sea and coastal environments. All isolates couple Fe oxidation to oxygen respiration, producing highly reactive Fe(III) oxyhydroxides that can adsorb or coprecipitate nutrients and metals. However, despite the biogeochemical importance of marine Fe oxidation, we are just beginning to learn how Fe-oxidizers function and influence marine ecosystems. With a recent surge of culturing and sequencing, there is now a significant set of data from which we can glean broader insights into microbial marine Fe oxidation.

The goal of this paper is to review our current knowledge of marine Fe-oxidizers through the lens of this increasingly well-established class of Fe-oxidizing bacteria (FeOB). We begin by describing the discovery of this novel class at an Fe-rich hydrothermal system at the Loihi Seamount in Hawaii. We lay out the evidence for microbially-driven Fe oxidation in this marine system, including new kinetics results from experiments with the Loihi isolate and model *Zetaproteobacteria Mariprofundus ferrooxydans* PV-1. Work at Loihi has inspired numerous studies of *Zetaproteobacteria* isolates, biominerals, and environmental distribution. In addition to reviewing these, we present a comprehensive reanalysis of *Zetaproteobacteria* diversity and ecology, enabled by the newly developed ZetaHunter classification program (2). We then use current genomic evidence to evaluate whether all members of this class have the potential to oxidize Fe and discuss inferred *Zetaproteobacteria* niches. Finally, we discuss our perspectives on open questions in *Zetaproteobacteria* evolution, ecology, and impacts on geochemical cycling. This article was submitted to an online preprint archive.

### *Zetaproteobacteria*: a novel class of marine Fe-oxidizing bacteria

The discovery of *Zetaproteobacteria* is a story that began decades before the class was proposed. The unusual morphology of biogenic Fe oxides have long been used to recognize microbial Fe oxidation in terrestrial environments. The twisted stalks of *Gallionella* and hollow sheaths containing cells of *Leptothrix* were described in terrestrial Fe-rich environments as early as the mid-1800s (3, 4). However, Winogradsky (5) was the first to confirm that *Leptothrix* required Fe(II) for growth, thus linking microbial activity with iron deposition in terrestrial environments (6). In the 1980s, similar structures were found in Fe-rich marine environments, including the Red Seamount of the East Pacific Rise, the Explorer Ridge, and Loihi Seamount (Figure S1) (7–11). This led to the assumption that these structures were made by *Gallionella* and *Leptothrix*, though these organisms were not detected in subsequent studies of marine Fe mats based on small subunit ribosomal RNA (16S) marker gene surveys (e.g. 12–14). Instead, Moyer et al. (12) discovered the first sequence of the novel *Zetaproteobacteria* class, though it wasn’t recognized at the time because there were no isolates or other closely related sequences. The first isolates, *Mariprofundus ferrooxydans* strains PV-1 and JV-1, were obtained from samples collected at Loihi Seamount near Hawaii in 1996-98 (1, 15). Additional surveys from Fe-rich environments provided related 16S gene sequences, which helped establish the *Zetaproteobacteria* as a monophyletic group within the Proteobacteria (1, 13, 16–18). The association of the *Zetaproteobacteria* and Fe-rich marine environments has been strengthened since these initial observations, with continued discovery of *Zetaproteobacteria* within Fe-rich saline environments.

### *Zetaproteobacteria* isolates: model systems for microbial Fe oxidation

The difficulty of culturing FeOB has been one of the main challenges in demonstrating microbial marine Fe oxidation. The first *Zetaproteobacteria* isolates were obtained using liquid and agarose-stabilized gradient tubes and plates designed to provide both Fe(II) and O_2_ in opposing gradients (15, 19). With this setup, Fe(II) is gradually released by dissolution of solid reduced Fe minerals (e.g. Fe(0) or FeS) at the bottom of the tube or plate while O_2_ diffuses from the headspace above. These culturing techniques make it difficult to control O_2_ and Fe(II) concentrations. To date, *Zetaproteobacteria* have not been culturable on solid media, so isolation requires serial dilution to extinction, with transfers every ∼2-3 days due to increasing autocatalytic Fe oxidation over time (20). In all, these challenges likely account for why so few *Zetaproteobacteria* have been isolated.

Despite these hurdles, *Zetaproteobacteria* representatives from two genera and eight separate OTUs have been successfully isolated (Table 1). These include seven isolates from Fe-rich hydrothermal Fe mats, and eight from coastal environments. *Mariprofundus ferrooxydans* PV-1 is the type strain of the most frequently isolated genus, and is an obligate neutrophilic autotrophic Fe-oxidizer. All but two other isolates are similarly obligate Fe-oxidizers. These two, *Ghiorsea bivora* TAG-1 and SV-108, are facultative Fe-oxidizers that are also capable of growth by H_2_ oxidation (21). Except for this instance, isolates vary primarily in their physiological preferences (Table 2), which are related to characteristics of their source environments.

**Table 1.**
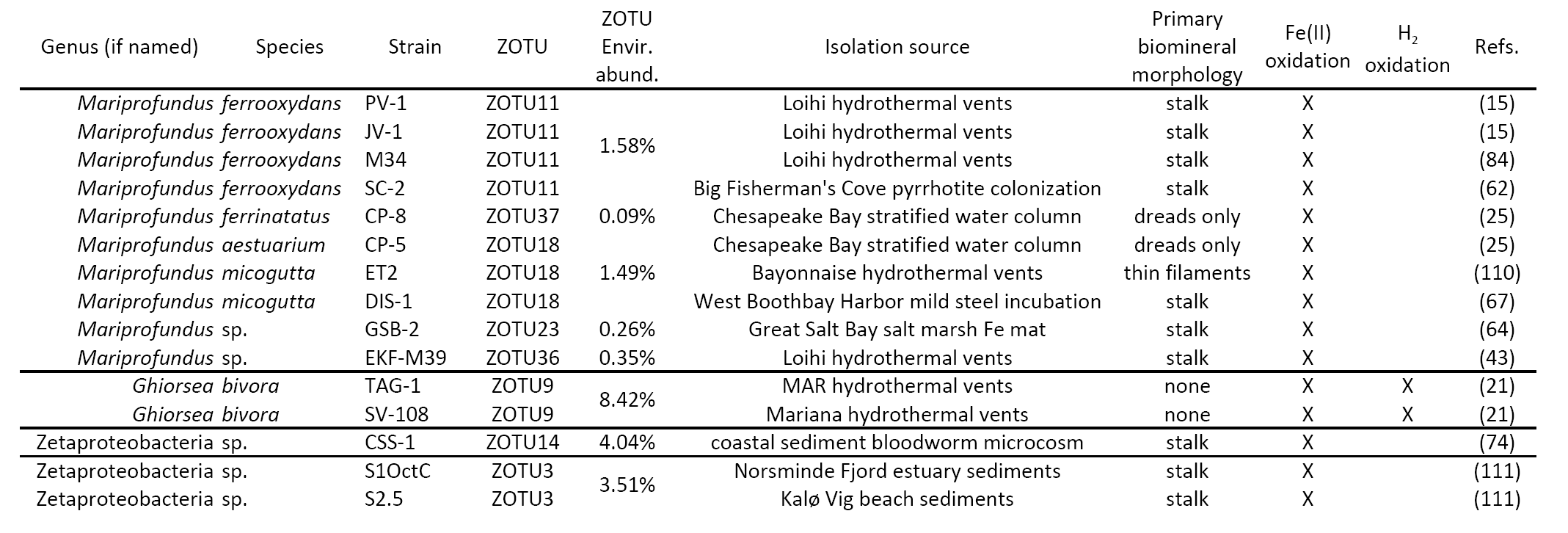
Isolates of the *Zetaproteobacteria* and their assigned ZOTUs, with representation in the environment, biomineral and metabolic properties, and references.

**Table 2.**
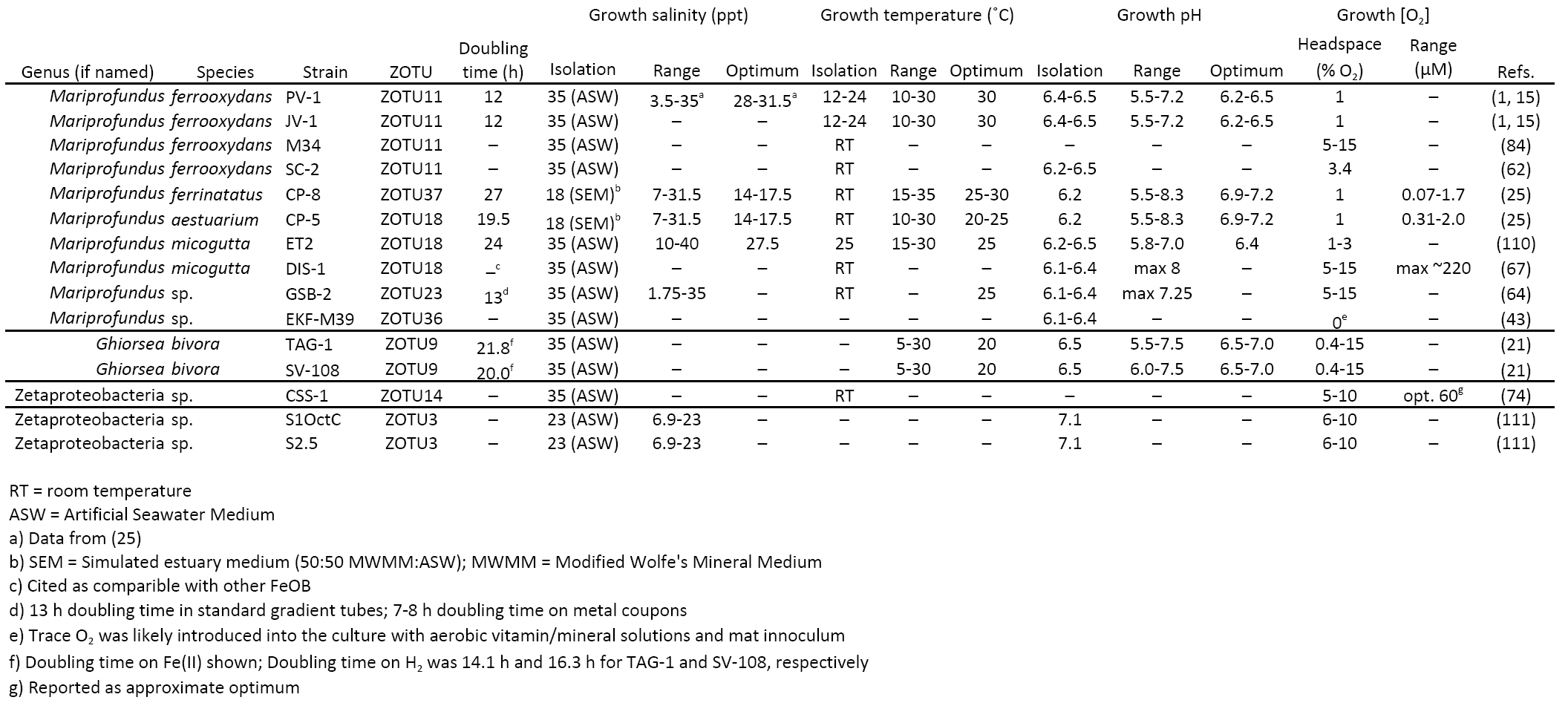
Growth preferences of *Zetaproteobacteria* isolates, including optimal growth salinity, temperature, pH, and oxygen concentrations.

*Zetaproteobacteria* isolates are generally microaerophiles originating from oxic-anoxic transition zones, where O_2_ concentrations are low, i.e. micromolar to submicromolar. Abiotic Fe oxidation is slow at these low O_2_ concentrations (22, 23), which allows the *Zetaproteobacteria* to compete. In terrestrial environments, kinetics experiments near 25°C suggest that biotic Fe oxidation is a significant component of total Fe oxidation below 50 μM and can outcompete abiotic Fe oxidation at 15 μM O_2_ (24). However, there is no kinetics data from marine FeOB. To understand the conditions where marine biotic Fe oxidation is competitive, we measured Fe oxidation kinetics using *M. ferrooxydans* PV-1 as a model (methods below). With this experiment, we have shown that PV-1 outpaces abiotic oxidation below 49 μM O_2_, and accounts for up to 99% of the Fe oxidation at 10 μM O_2_ (Figure 1; Table 3). In cultures of *M. aestuarium* CP-5 and *M. ferrinatatus* CP-8, oxygen concentrations ranged from 0.07-2.0 μM O_2_ within the cell growth band (25). This range of O_2_ growth conditions is well below the level at which almost all Fe oxidation was biotic for PV-1, suggesting that many *Zetaproteobacteria* are well adapted to compete and thrive under micromolar and submicromolar O_2_ concentrations. Such low O_2_ concentrations are common within the oxic-anoxic transition zones where the *Zetaproteobacteria* are found (e.g. 26, 27).

**Table 3.** Biotic and abiotic Fe oxidation rates of *Mariprofundus ferrooxydans* PV-1 under a range of O_2_ concentrations.

| O<sub>2</sub> conc.<br>μM | Fe(II) oxidation rate |  |  | % Biotic Fe(II)<br>oxidation | pH |
| --- | --- | --- | --- | --- | --- |
|  | biotic<br>μM Fe(II) hr<sup>-1</sup> | biotic<br>μM Fe(II) cell<sup>-1</sup> hr<sup>-1</sup> | abiotic<br>μM Fe(II) hr<sup>-1</sup> |  |  |
| 10.4 | 21.05 | 5.06E-04 | 0.32 | 98.5 | 6.63 |
| 10.4 | 24.69 | 5.53E-04 | 1.66 | 93.7 | 6.70 |
| 54.8 | 33.32 | 1.00E-03 | 39.43 | 45.8 | 6.67 |
| 79.7 | 5.57 | 1.25E-04 | 47.31 | 10.5 | 6.50 |

**Figure 1.**
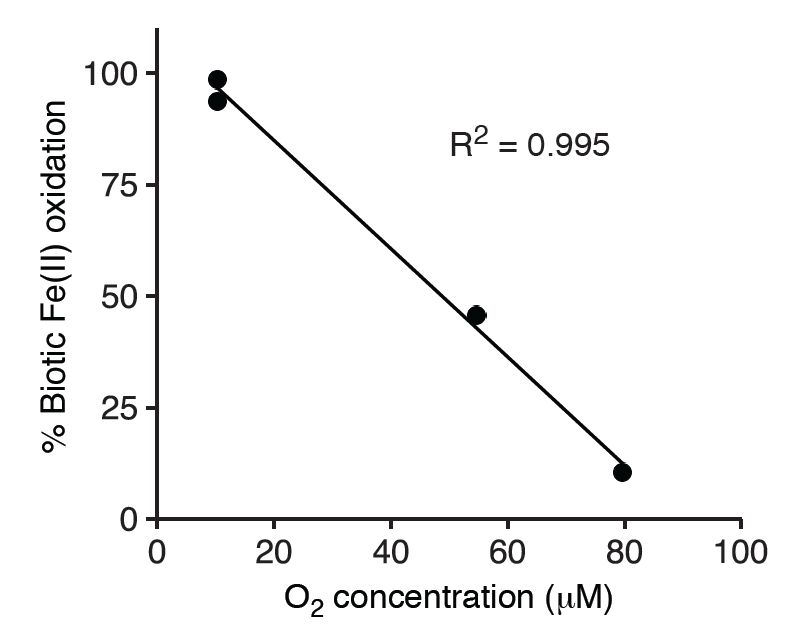
Relative rates of biotic versus abiotic Fe(II) oxidation at varying oxygen concentrations, using the model *Zetaproteobacteria* isolate, *M. ferrooxydans* PV-1. Range of 6.5 – 6.7 pH for experimental conditions. Further details in *Essential Methods*.

### Biomineral morphologies: form follows function

*M. ferrooxydans* PV-1 has been a model system for biomineralization by an obligate Fe-oxidizer. PV-1 cells form a twisted stalk (Figure 2A-C), so similar to the one formed by the terrestrial Fe-oxidizer *Gallionella ferruginea* that it could be mistaken for a *Gallionella* stalk (15). The stalk consists of individual filaments made of nanoparticulate Fe oxyhydroxides and acidic polysaccharides, controlling Fe mineral growth near the cell surface (28). Stalk growth was measured to be 2.2 μm h^-1^, or nearly 5x the width of a PV-1 cell per hour (28). The combination of this directed mineralization and a near-neutral cell surface charge explains how the cell remains remarkably free of encrustation (29). These encrustation avoidance mechanisms are important for any Fe-oxidizing microbe to avoid cell death by Fe mineral growth inside and outside the cell.

**Figure 2.**
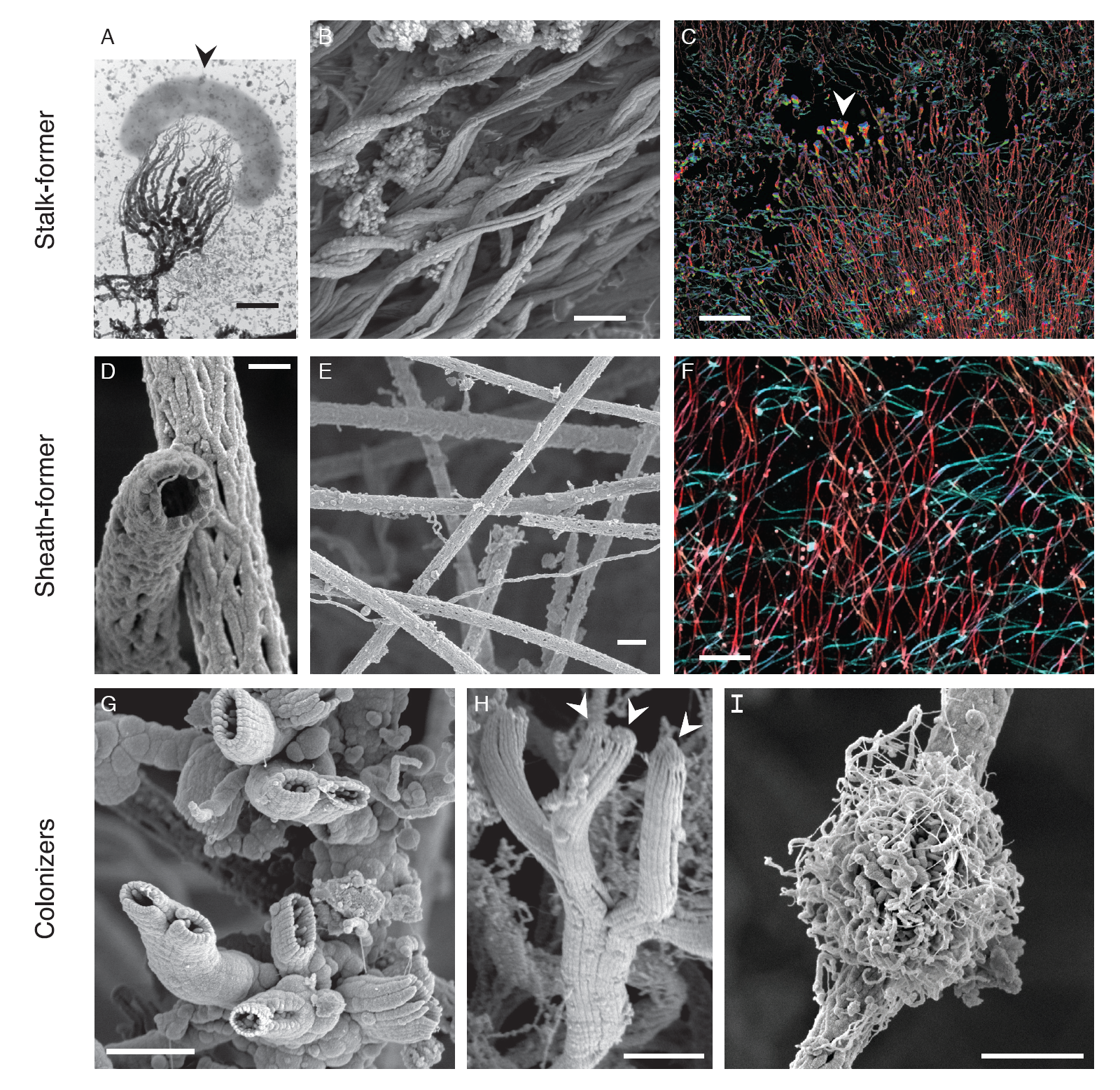
Morphologies of FeOOH biominerals known or suspected to be formed by the Fe-oxidizing *Zetaproteobacteria*. (A-C) Twisted stalks from individual cell to intact Loihi Fe microbial mats. (D-F) Sheaths from individual tubes to intact Loihi mats. (G-I) Fe biominerals attached to stalks and sheaths in Loihi mats. (A) A single *M. ferrooxydans* PV-1 cell (arrow) and its fibrillar stalk. (B) Intact curd-type mats (Figure 4A) are composed of parallel stalks. (C) Intact mat showing directional stalks that are interrupted (arrow), before biomineral production resumes. See Chan et al. (27) for details. (D) Hollow sheaths formed by the *Zetaproteobacteria*. (E) Intact veil-type mats (Figure 4B) are composed of sheaths. (F) Zoomed out intact mat with multiple sets of sheaths oriented in different directions (direction corresponds to color). (G-H) Short, Y-shaped tubular biominerals formed by *Zetaproteobacteria*. (H) Arrows show attached cells. (I) *Siderocapsa*-like nest-type biominerals; it is not known if they are formed by the *Zetaproteobacteria*. Scale bars: 0.5 μm (A,D), 2 μm (B,E,G-I), 100 μm (C,F). Images reproduced with permission: (A) from (28), (F, H) from (27). (B-I) from the samples described in (27). (D, I) imaged on JEOL-7200 field emission SEM.

While most *Zetaproteobacteria* isolates form a stalk, some make other biomineral morphologies (Table 1; Figure 3A). *Mariprofundus ferrinatatus* CP-8 and *M. aestuarium* CP-5 form shorter filaments that resemble the dreadlock hairstyle (Figure 3C) (25). Dreads were originally observed in terrestrial FeOB *Gallionellaceae Ferriphaselus spp.*, which makes both stalk and dreads (Figure 3B) (30). In both *Ferriphaselus* and the CP strains, the dreads are shed from cells. This suggests that dreads and similar structures are used specifically to avoid encrustation, whereas the stalk has other functions. PV-1 cells use the stalk as a holdfast to anchor the cell to surfaces (31). As the cell oxidizes Fe and produces new stalk the cell moves forward, leaving stalk behind. Since the stalk is rigid and anchored, this is a means of motility. Experiments in controlled Fe(II) and O_2_ gradients showed that PV-1 cells use their stalks to position themselves at an optimum position within that gradient, often forming filaments oriented toward higher O_2_ (31).

**Figure 3.**
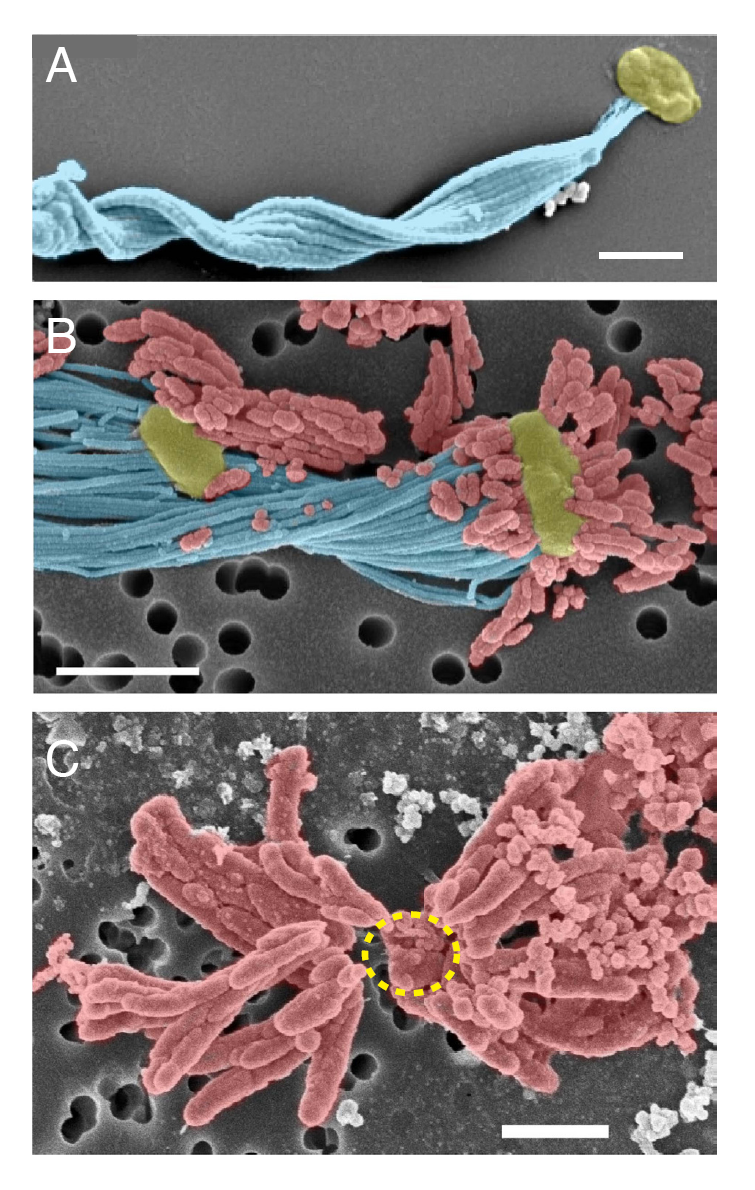
Colorized SEM images of stalk and dread biominerals of select marine and freshwater FeOB. (A) Twisted stalk (blue) formed by a *M. ferrooxydans* PV-1 cell (yellow). Most *Zetaproteobacteria* isolates form stalks. (C) In contrast, *M. aestuarium* CP-5 and *M. ferrinatatus* CP-8 produce only short dreads (red), which are easily shed from the cell (inferred cell position indivated by dashed yellow line). (B) Stalk (blue) and dreads (red) of the freshwater *Betaproteobacteria* FeOB *Ferriphaselus* sp. R-1 resembled the structures formed by marine FeOB. Scale bars = 1 μm. Images reproduced with permission: (A) from (27), (B) from (30), (C) from (25).

In the environment, such oriented filaments are common. At Loihi Seamount, curd-type mats (cohesive Fe mats with a bumpy surface reminiscent of cheese curds) often form directly above a vent orifice (Figure 4A) (27). Micrographs of intact curd mats showed centimeters-long, highly directional twisted stalks forming the mat architecture (Figure 2A-C). These stalks record the synchronous movements of a community of cells all growing and twisting in the same direction, as well as shifts in directionality in response to changes in the environment (27). The mechanism by which these cells actively control their directionality through stalk production is currently unknown.

**Figure 4.**
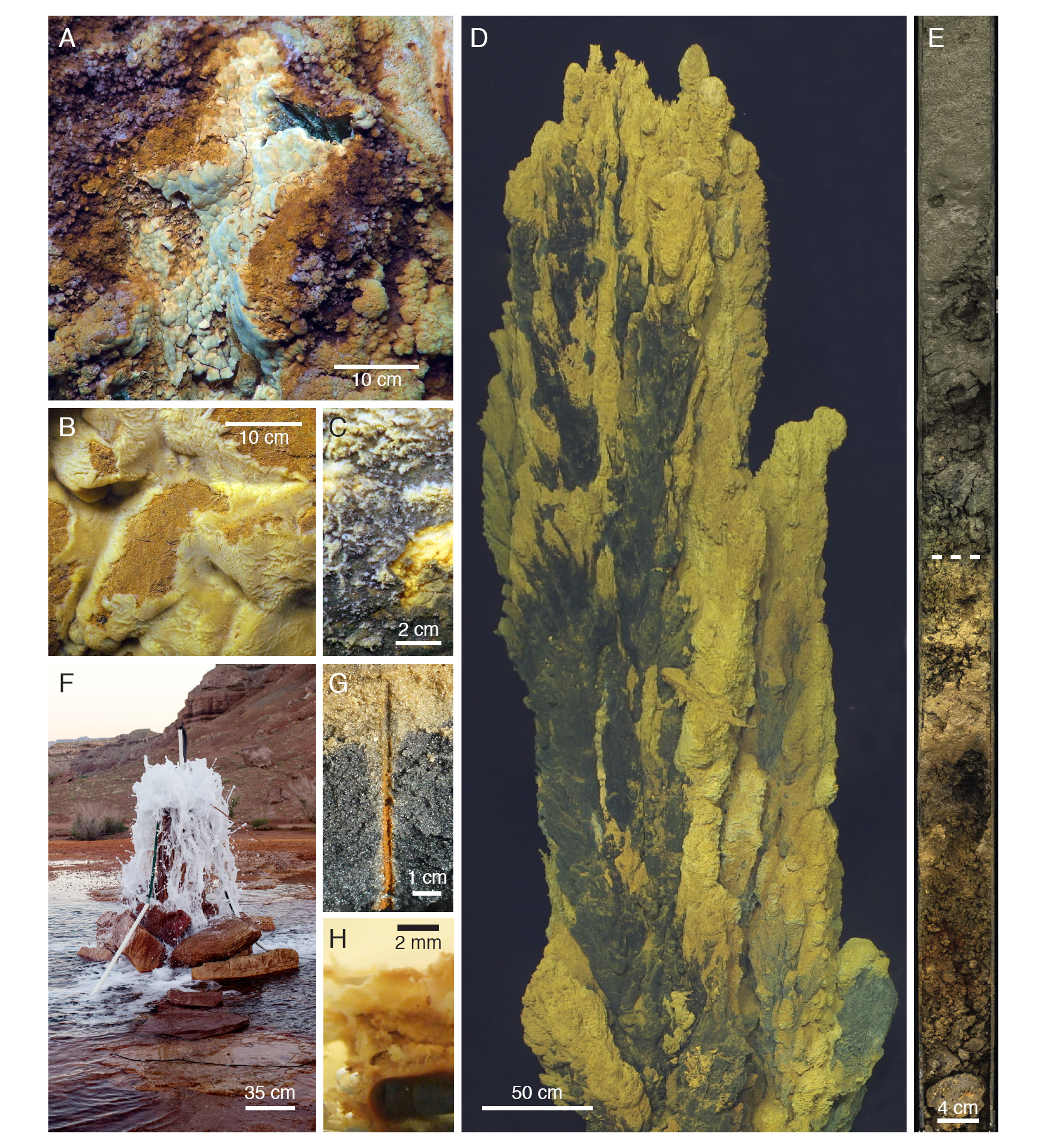
Photographs of *Zetaproteobacteria* habitats. (A-D) Marine hydrothermal vent mats, where *Zetaproteobacteria* have been found in highest abundance. (A) Curd-type and (B) veil-type Fe mats, from Loihi Seamount. (C) Mn-crusted Fe mat from the Ula Nui site, Loihi. Fe mat visible under broken surface (bottom right). (D) Fe mats on the Golden Horn Chimney, at the Urashima vent site, Mariana Trough. (E) Transition from reduced to Fe oxide-stained marine sediments (dashed line) in 26 mbsf core from the hydrothermal circulation cell of Iheya North vent field, Okinawa Trough. See (108) for details. (F) Terrestrial saline CO_2_-rich spring at Crystal Geyser, UT. (G) Fe oxide-coated worm burrows from the beach at Cape Shores, DE. (H) Mild steel corrosion biofilm formed by isolate *M.* sp. GSB-2. (H) Reproduced with permission from Joyce M. McBeth. (F) Original photography by Chris T. Brown; reproduced with permission.

Beyond stalks, Loihi Seamount also hosts sheath-rich veil-type mats, which form millimeters-thick Fe mat draped over rock or older Fe mat in diffuse venting environments (Figure 4B). These mats are created by organisms that form hollow Fe oxide sheaths (Figure 2D-F), similar to those produced by the terrestrial *Betaproteobacteria Leptothrix*. In the marine environment, however, these sheaths are formed by *Zetaproteobacteria*, informally called zetathrix (32). From studies based on the terrestrial *Leptothrix*, sheaths function similarly to stalks, with tens of cells producing the sheath and leaving it behind as the cells move forward (27). In Loihi Seamount intact veil mats, sheaths also leave a record of highly directional growth, despite oxygen profiles of these mats showing a shallow O_2_ gradient with O_2_ present throughout the mat (27). In both curd- and veil-type mats, *Zetaproteobacteria* work together to form a highly porous mat almost completely composed of biomineral filaments formed by cells, making these Fe mats different from other commonly studied mats or biofilms, which feature cells embedded in an exopolysaccharide matrix. The biomineral filaments forming the structure of the mat also frequently have Fe biominerals attached to them, suggesting FeOB also colonize the mat interior (Figure S1). Short branching hollow tubes, referred to as “y-guys”, are formed by the *Zetaproteobacteria* (Figure 2G,H) (33). However, the organisms forming other Fe biominerals have yet to be identified, including nest-like structures reminiscent of freshwater *Siderocapsa*-like organisms (Figure 2I) (34). The range of biomineral morphologies is related to differing biomineral functions, which likely correspond to different niches within the Fe mat habitat.

### Habitats of the *Zetaproteobacteria*

#### Loihi Seamount hydrothermal vents: a *Zetaproteobacteria*observatory

Most of what we know about *Zetaproteobacteria* is based on work at the Loihi Seamount (also spelled Lō’ihi Seamount), a Hawaiian submarine volcano, from long-term studies including the Iron Microbial Observatory (FeMO). Loihi Seamount is an ideal habitat for *Zetaproteobacteria*, with hydrothermal fluids rich in CO_2_ and Fe(II), and low in sulfide (<50 μM in vent fluids; undetectable in Fe mats) (10, 35, 36). Background seawater oxygen concentrations are ∼50 μM at the summit of Loihi Seamount, due to its location within the oxygen minimum zone (36). At the base of Loihi Seamount, the Ula Nui site has higher ambient O_2_ concentrations (145 μM), but lower venting temperatures (1.7°C average compared to ∼42°C average at the summit) (37). Low temperatures and low ambient O_2_ concentrations favor biotic Fe oxidation by reducing the abiotic rate at and below the mat surface (23, 38). Thus, the conditions at Loihi Seamount have favored the growth of microbial Fe mats ranging from centimeters to meters thick and up to 15 km^2^ (Figure 4A-C) (27, 37). The extensive Fe mats at Loihi Seamount may reflect years- to decades-long stable Fe mat production by the *Zetaproteobacteria*, based on productivity estimates (27, 33).

Loihi Seamount studies have provided the cornerstones of *Zetaproteobacteria* ecology. Since the discovery of *Zetaproteobacteria* in the 1990s (12, 15), five research expeditions from 2006-2013 have focused on *Zetaproteobacteria* succession, niche and species diversity, and genetic potential. Colonization experiments over 4-10 days showed that *Zetaproteobacteria* prefer low- to mid-temperature (from 22-60°C, avg 40°C) Loihi hydrothermal vents (14). This preference was reflected in longer term observations following the 1996 Loihi eruption, which showed *Zetaproteobacteria* increasing in abundance as high temperature vents cooled to pre-eruption temperatures and transitioned from sulfide-rich to Fe-rich fluids (36, 39–41). The bulk of the omic information on *Zetaproteobacteria* originates from Loihi Seamount, with the first isolate genome, single cell genomes, metagenome, and proteome all from Loihi sources, (42– 45).

#### Other hydrothermally-influenced habitats

Beyond Loihi, *Zetaproteobacteria* are hosted by many other hydrothermal systems. Extensive Fe mats form around vents at seamounts and island arc systems (Figure 4D) (41, 46–48). However, Fe mats have also been found at spreading ridge systems, within diffuse flow at the periphery of high-temperature chimneys and vents (49–52). Most hydrothermal Fe mats consist of biomineral morphologies similar to those at the Loihi Seamount (twisted stalks, sheaths, y-guys, etc.) (50, 51). However, mat textures and lithification can vary as a function of geochemistry (e.g. Mn, Si concentration) and rates of hydrothermal discharge (53, 54).

*Zetaproteobacteria* are also found in the marine subsurface. There, oxygenated seawater can mix with anoxic Fe-rich fluids, providing a favorable environment for Fe oxidation. *Zetaproteobacteria* have been observed up to 332 meters below the sea floor, within both hydrothermal recharge and cold oxic circulation cells (Figure 4E) (55– 57). In many near surface sediments, shallow mixing introduces O_2_ into an Fe-rich environment, leading to abundant *Zetaproteobacteria* populations (13, 18, 58). As hydrothermal systems age and cool, basalts and the minerals within inactive sulfide mounds can also serve as Fe(II) sources for *Zetaproteobacteria* (59–62). These studies show that the *Zetaproteobacteria* are abundant members of shallow and deep marine subsurface environments, one of the largest underexplored habitats in the oceans.

#### Coastal and terrestrial habitats

*Zetaproteobacteria* have only recently been discovered in coastal and terrestrial environments. Colonization experiments showed that *Zetaproteobacteria* biofilms grow on Fe(II) released from mild and carbon steel that is commonly used in ships and docks, suggesting that these FeOB contribute to corrosion (Figure 4H) (63–67). [For a review on the role of FeOB in biocorrosion, see Emerson (D. Emerson, submitted for publication; 67).] Fe(II) can also come from natural sources in coastal environments, originating from mineral weathering and transported in anoxic groundwater. Fe redox cycling at the oxic-anoxic transition zone of stratified estuaries can support the growth of *Zetaproteobacteria*, as evidenced by the isolation of *M. ferrinatatus* CP-8 and *M. aestuarium* CP-5 (25, 26). In near shore sediments, Fe-rich groundwater can support microbial communities with *Zetaproteobacteria* at the sediment surface (69–71). Also in these sediments, bioturbation from plant roots and animal burrows provides conduits of O_2_ to this Fe-rich groundwater. Biotic and abiotic Fe oxidation in these environments leads to the formation of Fe oxides, which coat sands, salt grass and mangrove roots, and burrows (Figure 4G) (64, 72–74). Beam et al. (74) found the abundance of *Zetaproteobacteria* within Fe oxide-coated worm burrows to be an order of magnitude higher than surrounding bulk sediment, suggesting that *Zetaproteobacteria* growth and biotic Fe oxidation can be favored in these bioturbated sediments. The Fe oxides produced in these environments can sequester toxins that adsorb to the mineral surface (75). Thus, FeOB activity could affect coastal water quality.

*Zetaproteobacteria* have generally been considered marine FeOB, detected at salinities up to 112 ppt in hypersaline brines (16, 76). Their occurrence in coastal environments provides the opportunity to delineate their minimum salinity requirements. McBeth et al. (77) surveyed Fe mats along the tidal Sheepscot River, Maine, as it entered the estuary, finding that *Zetaproteobacteria* only appeared in environments with 5 ppt salinity or higher. This explains why *Zetaproteobacteria* are not commonly found nor expected to be found in most terrestrial environments. This is further supported by deep sequencing of Fe-rich samples: our analyses of SRA data (Dataset S1) and Scott et al. (51) also determined that *Zetaproteobacteria* appear to be restricted to marine environments.

Thus, it was surprising and novel to find abundant populations of *Zetaproteobacteria* in CO_2_-rich terrestrial springs. Surveys of the 16S genes from carbonic springs at Tierra Amarilla Spring, New Mexico (∼9 ppt salinity) revealed a microbial population up to one third *Zetaproteobacteria* (78). Similarly, 16S and genomic work in CO_2_-rich Crystal Geyser, Utah, (∼11-14 ppt salinity; Figure 4F) found the *Zetaproteobacteria* to be both abundant and consistently present over a year of observation (79–81). These springs represent the first habitat with abundant populations of both *Zetaproteobacteria* and *Betaproteobacteria* FeOB (*Gallionellaceae*), whose abundance is likely driven by cycles of freshwater and saline subsurface groundwater mixing (81). The work at Crystal Geyser has produced full-length 16S sequences and the only terrestrial *Zetaproteobacteria* genomes (79–81). Our analysis of 16S phylogenetic placement and genomic clustering (by average nucleotide identity) suggests that *Zetaproteobacteria* populations in terrestrial subsurface environments are primarily novel and deeply branching *Zetaproteobacteria* (see further discussion of habitat selection and niche below).

#### Common habitat characteristics

In all, *Zetaproteobacteria* have been found in habitats sharing the following characteristics: 1) brackish to hypersaline water, 2) a supply of Fe(II), and 3) predominantly micro-oxic conditions. These conditions are widespread and found in diverse habitats, likely supplying multiple niches for the diversification and evolution of the *Zetaproteobacteria*.

### *Zetaproteobacteria* diversity

*Zetaproteobacteria* diversity has been defined using 16S gene *Zetaproteobacteria* operational taxonomic units (ZOTU; 97% similarity), based on sequences from isolates and environmental samples (Table 1; Dataset S1). Since their initial description, the *Zetaproteobacteria* class has remained a robust taxonomic group within the Proteobacteria (82, 83). A systematic analysis of 227 *Zetaproteobacteria* full-length 16S gene sequences yielded 59 ZOTUs (2), an increase from 28 ZOTUs in 2011 (84). The majority of these ZOTUs are contained within two families of the *Zetaproteobacteria*, based on sequence similarity (Figure 5A, Figure 6). Figure 5 shows key ZOTUs, which are frequently sampled and abundant in the environment and are primarily distinct monophyletic taxonomic groups by 16S (see detail in Figure S2). These 15 ZOTUs represent 83% of sequences found in the environment. ZOTUs 1 and 14 are the one exception to monophyly by 16S, yet do form distinct monophyletic groups in a concatenated tree of 12 ribosomal proteins (Figure 6; phylogenetic methods in *Supplemental Methods*). ZOTUs that include isolates represent only 20% of environmental sequences (Table 1), showing that the *Zetaproteobacteria* are largely uncultivated.

**Figure 5.**
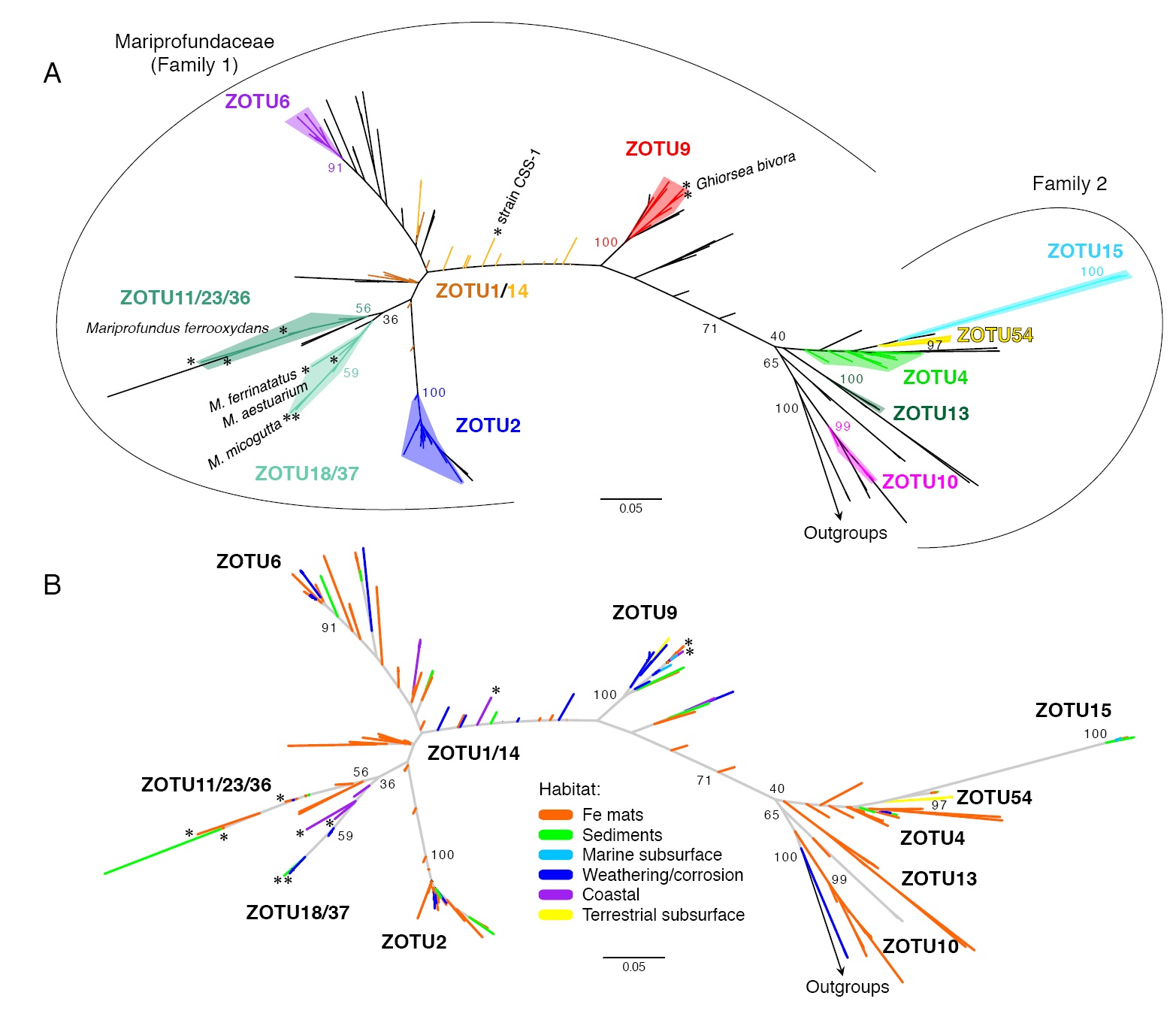
Maximum likelihood phylogenetic tree showing *Zetaproteobacteria* 16S gene diversity, (A) colored by ZOTU and (B) colored by habitat type where the sequences were sampled. A total of 59 ZOTUs have been classified, though only the most frequently sampled are shown in the figure above. ZOTUs 1 and 14 are poorly resolved by the 16S gene. Published isolates of the *Zetaproteobacteria* are starred and labeled. Phylogenetic trees were colored automatically using Iroki (109).

**Figure 6.**
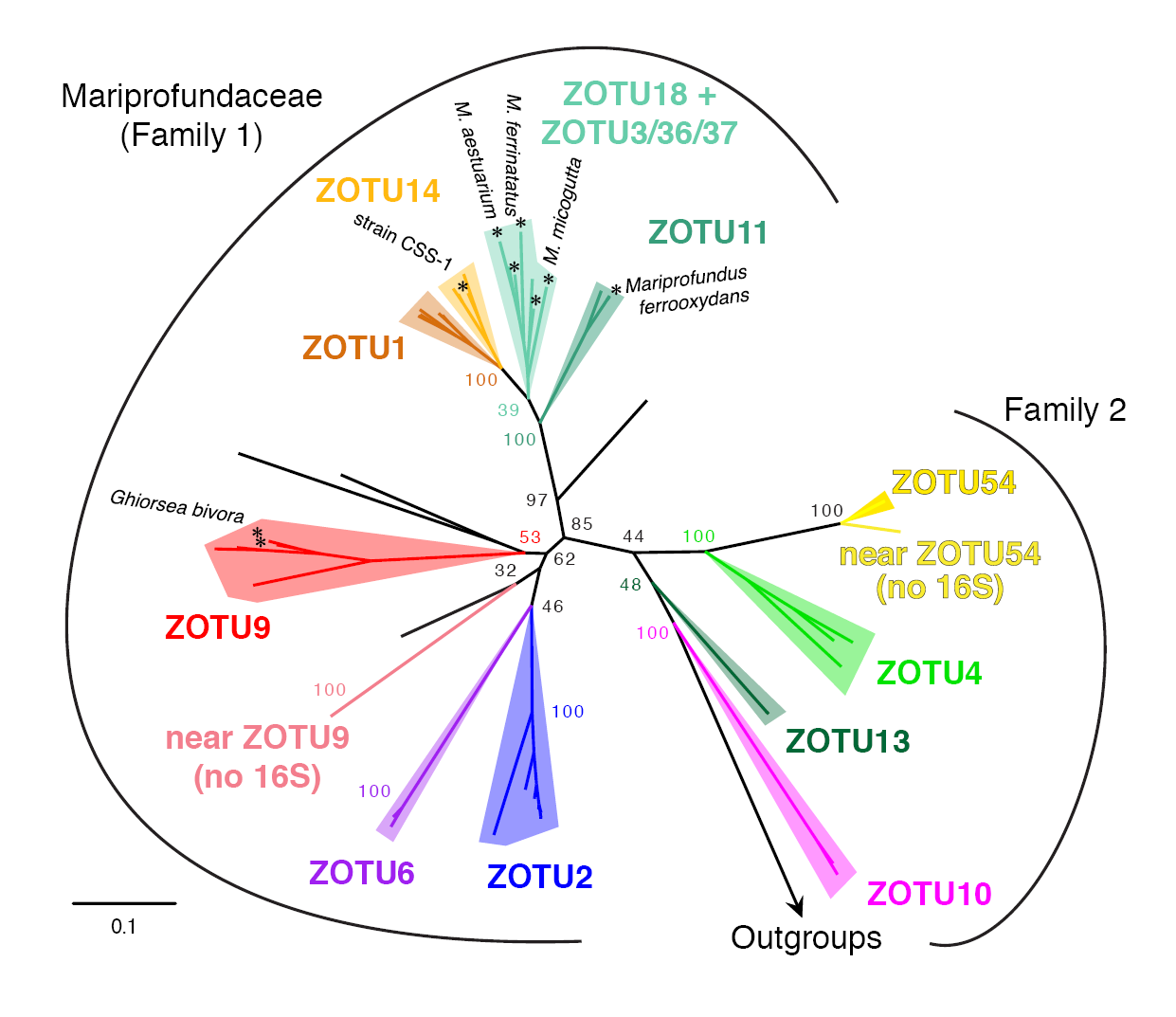
Maximum likelihood phylogenetic tree showing a more robust phylogenetic placement of ZOTUs based on the concatenated alignments of 12 ribosomal proteins. Data from isolate, SAG, and MAG genomes (see *Supplemental Methods*). In this tree, ZOTUs 1 and 14 are monophyletic entities. Taxa near ZOTU9 and near ZOTU54 are only represented by genomes; these clades have only been found in the terrestrial CO_2_-rich spring waters of Crystal Geyser, UT.

In order to compare *Zetaproteobacteria* diversity across studies, we used ZetaHunter, a classification pipeline designed to rapidly and reproducibly classify ZOTUs (2). We classified publicly available *Zetaproteobacteria* full- and partial-length 16S gene sequences from SILVA, IMG, and NCBI SRA (total of 1.2 million sequences from 93 studies; summary of samples in Dataset S1; methods in *Supplemental Methods*) (85–88). This work provided the basis for the habitat and diversity analysis below, while also allowing us to correct previous ZOTU assignments (Table S1).

#### Connecting *Zetaproteobacteria* diversity, habitat, and niche

*Zetaproteobacteria* diversity is driven primarily by the variety of niches they inhabit. A niche is the set of conditions favorable for growth, which are further influenced, or partitioned, by inter- and intra-species population dynamics in the environment (89). A challenge in microbial ecology is to tease apart the niche of an organism through sampling at the appropriate resolution. For most marine environments, we lack the highly resolved chemical and spatial information to describe niches. However, we can look for patterns of associations between different *Zetaproteobacteria* and their habitats to understand where and the extent to which *Zetaproteobacteria* niches may overlap.

Each habitat displayed distinct and abundant ZOTUs, indicating that habitats can host a specific set of niches that support these ZOTUs (Table 4; Dataset S1). In particular, dominant ZOTUs differ between habitat types, suggesting that each habitat has a set of dominant niches that favor the growth of those particular ZOTUs. ZOTU54 is a striking example of a dominant ZOTU clearly successful within the terrestrial subsurface fluid environment. ZOTU54 is a deep-branching ZOTU that is primarily limited to this environment (78–81). The distribution of ZOTU54 suggests that it is endemic or adapted to thrive within terrestrial carbonic Fe-rich springs. However, while it is frequently found at high abundance in terrestrial springs, ZOTU54 is also found in other habitats at very low abundance, including hydrothermal Fe mats (Dataset S1). In fact, many ZOTUs span habitats (Figure 5B; Dataset S1), suggesting that similar niches supporting these ZOTUs can exist in multiple habitats.

**Table 4.**
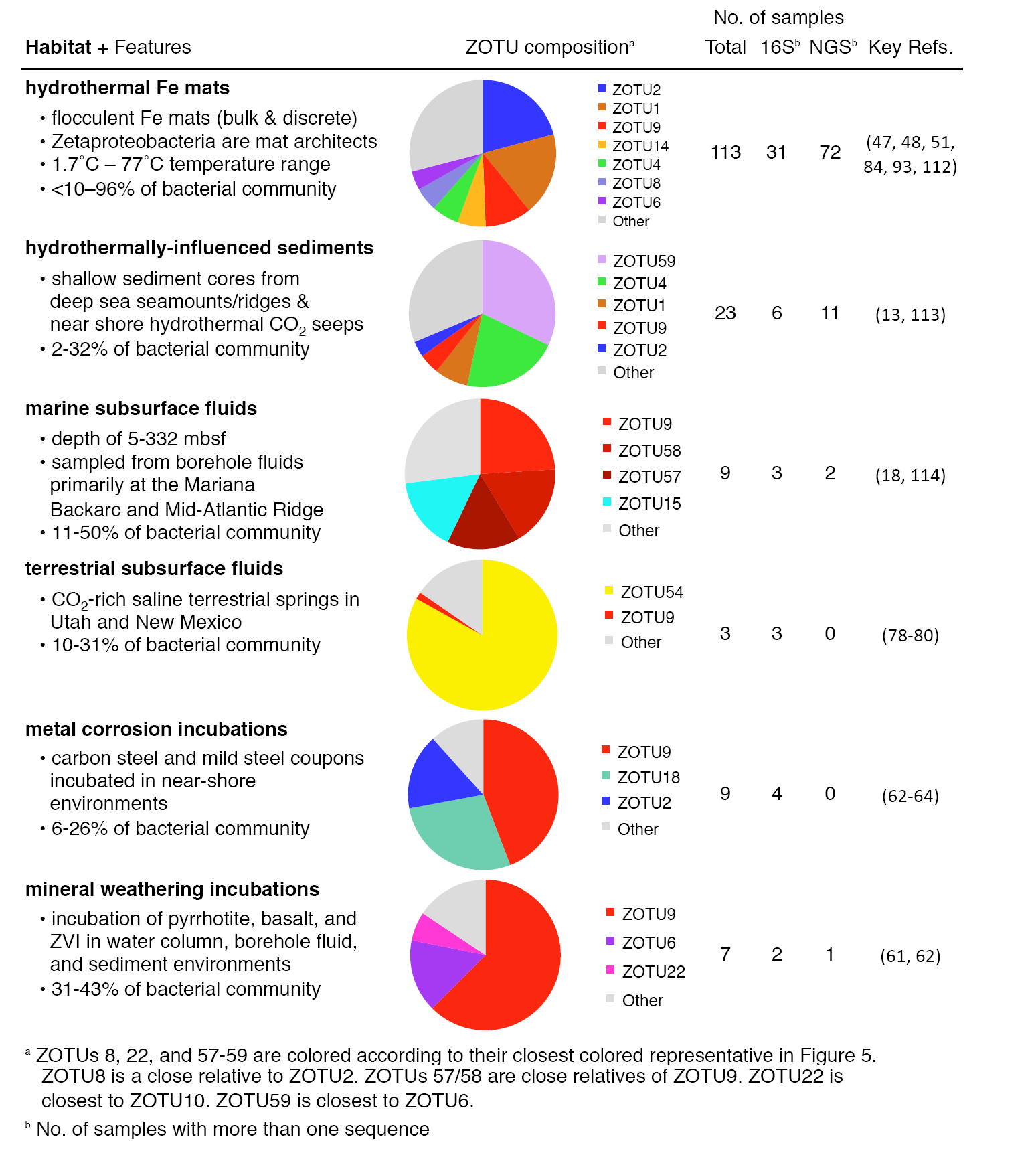
Summary of the habitats where the *Zetaproteobacteria* are found in high abundance using data from Dataset S1.

| Habitat + Features | ZOTU composition<sup>a</sup> | No. of samples |  |  | Key Refs. |
| --- | --- | --- | --- | --- | --- |
|  |  | Total | 16S<sup>b</sup> | NGS<sup>b</sup> |  |
| **hydrothermal Fe mats**<br>• flocculent Fe mats (bulk & discrete)<br>• Zetaproteobacteria are mat architects<br>• 1.7°C – 77°C temperature range<br>• <10–96% of bacterial community | ■ ZOTU2<br>■ ZOTU1<br>■ ZOTU9<br>■ ZOTU14<br>■ ZOTU4<br>■ ZOTU8<br>■ ZOTU6<br>■ Other | 113 | 31 | 72 | (47, 48, 51, 84, 93, 112) |
| **hydrothermally-influenced sediments**<br>• shallow sediment cores from deep sea seamounts/ridges & near shore hydrothermal CO<sub>2</sub> seeps<br>• 2-32% of bacterial community | ■ ZOTU59<br>■ ZOTU4<br>■ ZOTU1<br>■ ZOTU9<br>■ ZOTU2<br>■ Other | 23 | 6 | 11 | (13, 113) |
| **marine subsurface fluids**<br>• depth of 5-332 mbsf<br>• sampled from borehole fluids primarily at the Mariana Backarc and Mid-Atlantic Ridge<br>• 11-50% of bacterial community | ■ ZOTU9<br>■ ZOTU58<br>■ ZOTU57<br>■ ZOTU15<br>■ Other | 9 | 3 | 2 | (18, 114) |
| **terrestrial subsurface fluids**<br>• CO<sub>2</sub>-rich saline terrestrial springs in Utah and New Mexico<br>• 10-31% of bacterial community | ■ ZOTU54<br>■ ZOTU9<br>■ Other | 3 | 3 | 0 | (78-80) |
| **metal corrosion incubations**<br>• carbon steel and mild steel coupons incubated in near-shore environments<br>• 6-26% of bacterial community | ■ ZOTU9<br>■ ZOTU18<br>■ ZOTU2<br>■ Other | 9 | 4 | 0 | (62-64) |
| **mineral weathering incubations**<br>• incubation of pyrrhotite, basalt, and ZVI in water column, borehole fluid, and sediment environments<br>• 31-43% of bacterial community | ■ ZOTU9<br>■ ZOTU6<br>■ ZOTU22<br>■ Other | 7 | 2 | 1 | (61, 62) |

<sup>a</sup> ZOTUs 8, 22, and 57-59 are colored according to their closest colored representative in Figure 5. ZOTU8 is a close relative to ZOTU2. ZOTUs 57/58 are close relatives of ZOTU9. ZOTU22 is closest to ZOTU10. ZOTU59 is closest to ZOTU6.
<sup>b</sup> No. of samples with more than one sequence

Next, we looked for patterns in ZOTU associations with each other, mapping connections in a ZOTU network (Figure 7). This network shows which ZOTUs are found in isolation and which co-occur in the same sample, with connection thickness showing the strength of those connections. Further, the network layout is based on the frequency of co-occurrence, so when two ZOTUs commonly co-occur, they are closer together and connected by thicker lines. Most ZOTUs co-occur with others (Figure 7), and these connections are not random. Some ZOTUs co-occur more frequently, forming clusters of interconnected ZOTU nodes (Clusters 1-3, Figure 7). The most abundantly sampled ZOTUs form a central cluster (Cluster 2), sharing a common set of niches most frequently sampled in the hydrothermal Fe mat environment. Cluster 3 centers around ZOTUs found together in hydrothermal Fe mat samples from the Mariana Arc, but which are not common in other environments. Cluster 1 is dominated by ZOTUs from samples associated with metal corrosion and mineral weathering, which suggests that these habitats host niches distinct from those in hydrothermal Fe mat habitats. Overall, these clusters highlight ZOTUs with niches that frequently overlap, suggesting those niches, and thus the growth requirements of these ZOTUs, are compatible.

**Figure 7.**
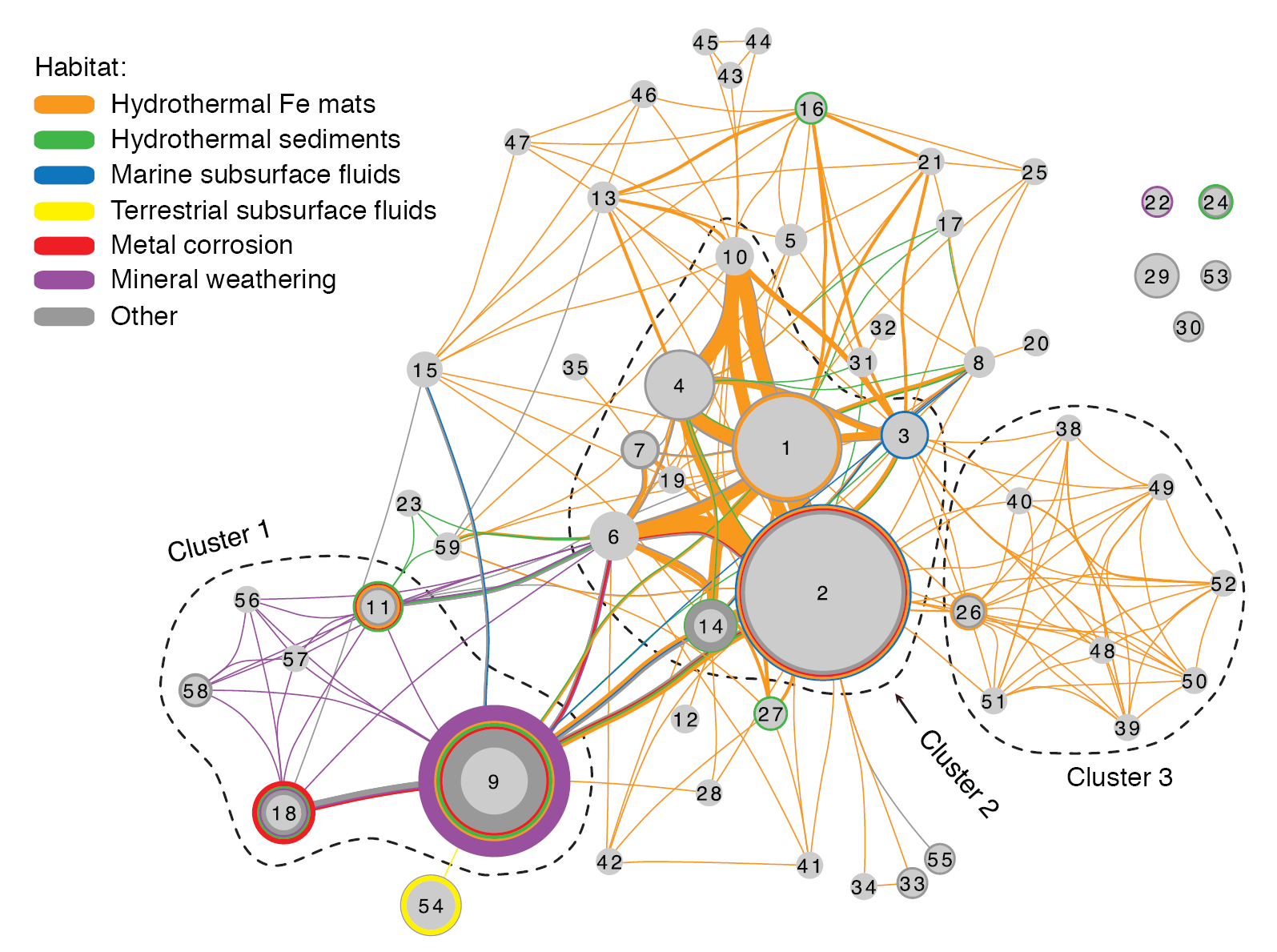
*Zetaproteobacteria* OTU network showing the association of known ZOTUs within the specified habitats. Lines connect ZOTUs that are found in the same sample, with thickness representing the frequency of that association in multiple samples. Colored circles surrounding ZOTU nodes show samples where only a single ZOTU was found. ZOTU nodes are sized according to their environmental abundance. Placement of ZOTUs in the network was determined automatically, based on the frequency of co-occurrence (Cytoscape’s edge-weighted, spring embedded layout). Dotted lines denote ZOTU clusters common to the following habitats: Cluster 1, metal corrosion and mineral weathering; Cluster 2, ubiquitous Fe mats; Cluster 3, Mariana Trough Fe mats. Isolate sequences are not shown. Dataset based on SILVA release 128.

A combination of ZOTU environmental distribution, habitat characteristics, and isolate physiology can help us understand niche. Here, we use this approach to describe ZOTU9, as an example. This ZOTU is a key player in marine subsurface fluids, metal corrosion, and mineral weathering habitats (see Cluster 1, Figure 7; Table 4). In mineral weathering habitats, ZOTU9 is frequently the only ZOTU (visualized as a colored circle around the ZOTU node; Figure 7). For example, ZOTU9 was the only *Zetaproteobacteria* detected within a basaltic glass weathering enrichment, making up 39% of the bacterial community by 16S sequencing estimates (61). These habitat associations suggest that growth of ZOTU9 is favored when metal corrosion or mineral weathering is a source for Fe(II). This association likely relates to these sources producing H_2_ as well as Fe(II). Metals composed of zero valent Fe produce H_2_ as a byproduct of anaerobic corrosion (90). Fe minerals can be a source of H_2_ because they catalyze hydrolysis and induce radiolysis of water (91, 92). The production of H_2_ is known to benefit some members of ZOTU9, including the Fe- and H_2_-oxidizing isolates *Ghiorsea bivora* TAG-1 and SV-108 (21). The genetic machinery required for H_2_ oxidation has also been found in two ZOTU9 single amplified genomes (43, 51), which suggests that H_2_ oxidation may be common in other members of ZOTU9. From these observations, we conclude that both Fe(II) and H_2_ play a central role in the niche of ZOTU9 and the habitats where it can be found. By combining isolate physiology with habitat distribution patterns, we can identify key features of a ZOTU’s niche.

#### Spatial and taxonomic resolution in *Zetaproteobacteria* ecology

Hydrothermal Fe mats have opposing gradients of Fe(II) and O_2_ and a complex internal structure (e.g. Figure S1) (27, 36). This heterogeneity leads to multiple niches at small spatial scales, suggesting that high-resolution sampling could help us better understand ZOTU niches in this habitat. Initial bulk techniques for sampling collected liters of mat material, and a single sample could contain all major ZOTUs (84). Therefore, new collection devices were engineered to sample small volumes (50-75 mL) at centimeter spatial resolution (50). From these discretely sampled Fe mats, we increased the resolution of our ZOTU network (Figure S3). Of the 29 ZOTUs found within Fe mat habitats, 17 showed a preference for a specific Fe mat type. However, the high connectivity of abundant ZOTUs remained, even when considering more highly resolved sampling. This result suggests that these ZOTUs share compatible niches at the centimeter scale in Fe mat habitats.

Taxonomic resolution can also affect our understanding of *Zetaproteobacteria* ecology through the lumping or splitting of ecologically-distinct groups. The ZOTU classification may in certain cases be too coarse, representing multiple related populations that have different niches. For example, Scott et al. (93) found multiple oligotypes (ecological units defined by informative sequence variability) within each ZOTU. While multiple oligotypes do not necessarily suggest each has a distinct niche, for ZOTU6, only one oligotype differed in abundance over a transect approaching the hydrothermal vent orifice. This abundance change suggested that a subpopulation of ZOTU6 prefers higher flow conditions, warmer temperatures, and/or the differing geochemistry found near the vent (93). Results like this warrant a more resolved *Zetaproteobacteria* taxonomy, which could be aided by whole genome comparisons.

### Using genomics to understand metabolic potential and niche

#### Carbon fixation

All *Zetaproteobacteria* isolates are obligate autotrophs, using the Calvin-Benson-Bassham (CBB) cycle to fix carbon. Similarly, all ZOTUs sampled to date have the ribulose-1,5-bisphosphate carboxylase oxygenase gene (RuBisCO; key enzyme in the CBB cycle), suggesting carbon fixation by this pathway is a shared capability across the class. The isolates of *Mariprofundus ferrooxydans,* strains PV-1, JV-1, and M34, all encode the genes for both Form I (O_2_-insensitive) and Form II (O_2_-sensitive) RuBisCO (42, 94). Similarly, both forms are encoded by *Mariprofundus micogutta* DIS-1, which was specifically isolated to be more aerotolerant (67). However, most *Zetaproteobacteria* SAG and MAG genomes only encode Form II RuBisCO (43, 80, 95), suggesting most Zetaproteobacteria are specifically adapted to lower O_2_ concentrations.

#### Energy metabolism: Are all *Zetaproteobacteria* Fe-oxidizers?

The *Zetaproteobacteria* are often associated with high Fe environments, and all isolates of the *Zetaproteobacteria* are capable of Fe oxidation. These observations have led to the current assumption that all *Zetaproteobacteria* are capable of Fe oxidation. To test this assumption, we first have to understand the mechanism of Fe oxidation in the marine environment.

Initial genome analysis of PV-1 led to the proposal of the alternative complex III (ACIII) as part of an iron oxidase complex (42). Follow up studies later changed this model, suggesting ACIII was involved in reverse electron transport (30, 44, 45). However, Field et al. (43) and Chiu et al. (25) isolated *Zetaproteobacteria* isolates that lacked ACIII but were still capable of Fe oxidation. Furthermore, only 2 of 23 *Zetaproteobacteria* SAGs have the ACIII gene, and these 23 SAGs represent the majority of *Zetaproteobacteria* diversity (43). Combined, this evidence showed that ACIII is not a critical component of the Fe oxidation pathway.

The putative Fe oxidase, Cyc2, and another cytochrome Cyc1 were first identified in *Zetaproteobacteria* by Barco et al. (45) through a proteome analysis of PV-They were initially identified through comparison of the proteome with the closely related *M ferrooxydans* M34 genome. Their presence in the proteome suggested that the *cyc1* and *cyc2* genes were missing from the PV-1 draft genome due to gaps in the assembly, which was confirmed by resequencing (45). The Cyc2 protein from PV-1 is a homolog of the biochemically-characterized Cyc2 Fe oxidase from *Acidithiobacillus ferrooxidans* (22% amino acid identity) (96). Based on this, the Fe oxidation pathway model for the *Zetaproteobacteria* was revised (Figure 8). Cyc2 homologs have been found in other *Zetaproteobacteria* and other neutrophilic FeOB, strengthening the proposed pathway (97, 98). In fact, every single ZOTU that has a genomic representative, including ZOTUs without isolates, has a homolog of this putative Fe oxidation gene, consistent with the notion that all *Zetaproteobacteria* are Fe-oxidizers (43, 95).

**Figure 8.**
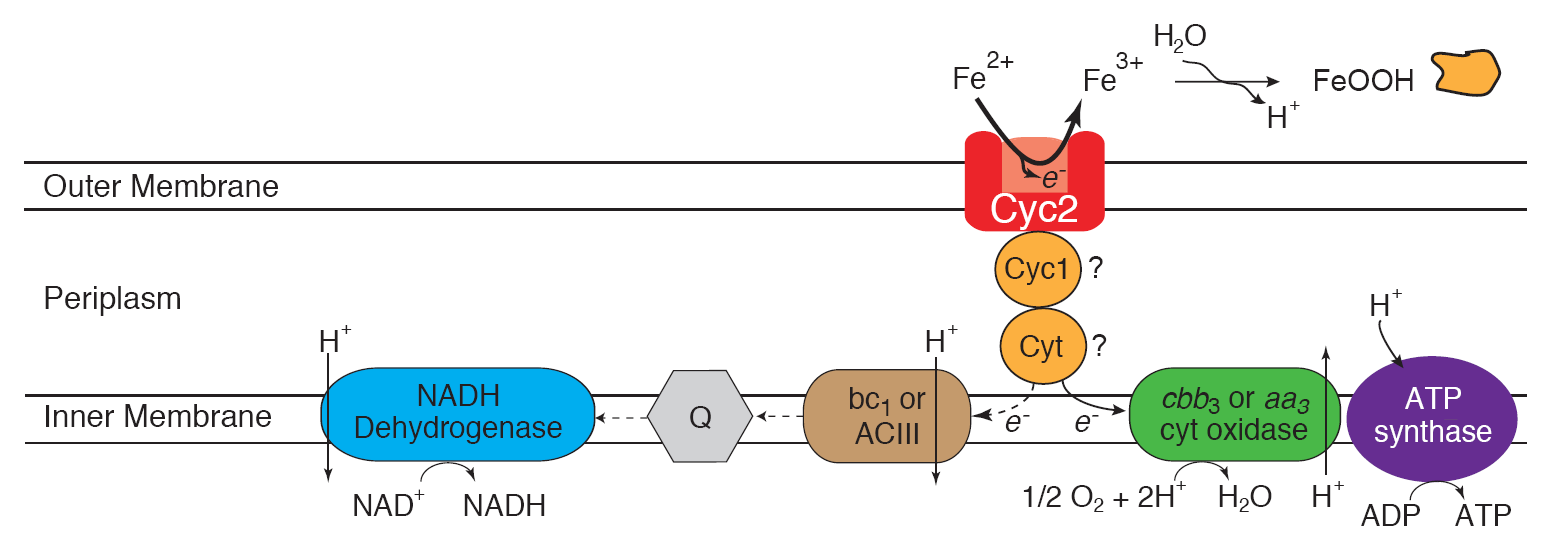
Model for Fe oxidation in the *Zetaproteobacteria* modified from (45). An electron from Fe(II) is passed from Cyc2 to a periplasmic electron carrier (Cyc1 and/or other c-type cytochrome) before being passed to the terminal oxidase (cbb_3_- or aa_3_-type cytochrome c oxidases), generating a proton motive force. For reverse electron transport, the electron from the periplasmic carrier is passed to the bc_1_ complex or alternative complex III before being passed to the quinone pool where it is used to regenerate NADH.

#### Genomic clues to niche based on O_2_ and nitrogen

Genomic evidence suggests that adaptation to differing O_2_ conditions plays a role in ZOTU niches. Three terminal oxidases have been found within *Zetaproteobacteria* genomes: cbb_3_- and aa_3_-type cytochrome c oxidases and the cytochrome bd-I ubiquinol oxidase. These have different affinities for oxygen, which would influence the niche of each ZOTU; K_m_’s of 230-300 nM (cbb_3_, bd-I) to 4.3 μM (aa_3_) are reported (99, 100). The cbb_3_-type terminal oxidase gene is found in most of the *Zetaproteobacteria*, sometimes in multiple copies, suggesting a predominant preference for very low O_2_ concentrations (submicromolar) (43). However, the complete genomes of *M. aestuarium* and *M. ferrinatatus* contain only the higher-O_2_ adapted aa_3_-type terminal oxidase gene, which helps explain their adaptation to frequently higher O_2_ concentrations of their tidally-mixed water column habitat (25). Many *Zetaproteobacteria* genomes have multiple terminal oxidases, suggesting they are adapted to fluctuating oxygen conditions (43, 95). ZOTU10 and the isolate *M. micogutta* DIS-1 may have a higher tolerance for such conditions with increased numbers of genes for O_2_ radical protection (43, 67).

The genetic potential for nitrogen transformations differentiates marine and terrestrial *Zetaproteobacteria*. In the marine environment, most ZOTUs are capable of assimilatory nitrate reduction to ammonium (*nasA, nirBD*) (43, 95). In terrestrial Fe-rich springs such as Crystal Geyser, *Zetaproteobacteria* genomes lack these genes, but many possess nitrogen fixation genes (e.g. *nifH*) (79, 80). In contrast, only three marine isolates (*Mariprofundus* strains DIS-1, EKF-M39, and M34) and one MAG outside of *Mariprofundus* possesses *nif* genes (43, 67, 95), and, as yet, it has not been experimentally shown that these isolates fix N_2_. Supporting these genomic observations, the *nifH* gene is rarely detected in marine Fe mats (101). From these patterns, the differences between these *Zetaproteobacteria* likely correspond with differences in nitrate and ammonium availability in these habitats; nitrate is below detection at Crystal Geyser compared to concentrations up to 32 μM within Loihi Fe mats (79, 102). Nitrogen transformations and O_2_ tolerance obviously play a role in many *Zetaproteobacteria* niches, but there are likely other conditions driving niche diversity yet to be discovered.

#### Outstanding questions and opportunities

Over the last two decades, *Zetaproteobacteria* have been established as a diverse, taxonomically-robust class, which thrive in a wide range of Fe-rich habitats. Environmental studies, isolate experiments, and genomic analyses have given insight into how they use biomineralization and metabolic strategies to succeed. Building on this work, we are poised to address a number of intriguing questions.

##### How did the *Zetaproteobacteria* come to specialize in Fe oxidation?

Thus far, genomic evidence suggests that all *Zetaproteobacteria* are Fe-oxidizers. If this is true, the *Zetaproteobacteria* would be an interesting model system in which to explore the selection and evolution of a particular metabolic specialty. The answer to this question likely rests on the complex challenges of Fe oxidation at circumneutral pH. *Zetaproteobacteria* must position themselves at specific environmental interfaces to gain energy from Fe oxidation. Meanwhile, they must compete with or tolerate abiotic reactions of Fe(II) with O_2_ and nitrogen compounds, which can form O_2_ radicals and toxic nitric and nitrous oxides (103, 104). They produce intricate biomineral structures, which allow them to avoid encrustation, control motility, and construct mats. Thus, microbial Fe oxidation appears to be a complex physiological trait, which is much more likely to be inherited vertically through descent rather than transmitted horizontally (105). Since Fe oxidation is a complex trait, the *Zetaproteobacteria* probably acquired the Fe oxidation capability before the group diverged. It is unclear where the Fe oxidation trait originated, but as we determine the genetic basis for the trait and adaptations to its challenges, comparative genomics of various FeOB will allow us to understand these evolutionary relationships.

##### What are the drivers of *Zetaproteobacteria* diversification?

The *Zetaproteobacteria* have diversified into at least 59 operational taxonomic units, which we can now track using ZetaHunter (2). Given the increasing number of available genomes, the next logical step is to develop a systematic taxonomy based on both 16S gene and phylogenomics analysis. Ultimately, diversification is driven by the range in environmental niches. We will improve our understanding as we continue to study environmental distribution, physiology of new isolates, and genomes, especially as we focus our explorations beyond the well-studied hydrothermal vents. We may be able to define niches better via discrete sampling, though there are practical lower limits to sample size and spatial resolution. Although intact samples are challenging to obtain, the effort is worthwhile in order to use imaging-based techniques (e.g. FISH), coupled to chemistry and activity measurements (e.g. elemental mapping, SIP) to discern millimeter- and micron-scale associations. We are just beginning to discover the variety of adaptations across genomes. As genome analyses progress, patterns of functional genes and phylogeny will elucidate the drivers of *Zetaproteobacteria* diversification. In turn, genomic clues can help us culture novel organisms, which will be key to demonstrating particular roles. The integrated results of these studies will show how these organisms have evolved to occupy particular niches, and how they could work together to influence the geochemistry of Fe-rich habitats.

##### How do *Zetaproteobacteria* affect geochemical cycling, and how can we track these effects?

Now that we know the basics of *Zetaproteobacteria* metabolisms and potential geochemical effects, we can move toward detecting this influence in the environment and determining the controls on those effects. The key will be developing ways to track *Zetaproteobacteria* activity, and relating this to quantitative effects. There is no clear biotic Fe isotopic signature that can be used to assess the activity of microbially-mediated Fe oxidation (106). An alternative is to track activity via gene expression. Traditionally, this would be done via a marker gene for Fe oxidation. The *cyc2* gene may work if its expression proves to be specific to Fe oxidation. However, now with (meta)transcriptomic approaches, we can use multiple genes (e.g. the whole Fe oxidation pathway, linked with C fixation and other pathways). With the *Zetaproteobacteria*, this will be an iterative exercise, as we are still determining/validating the genes involved in Fe oxidation and other metabolisms. This will be most straightforward in *Zetaproteobacteria*-dominated hydrothermal Fe mat environments, but work in other environments will improve our understanding of the range of their effects on geochemical cycling. As *Zetaproteobacteria* are widespread in diverse environments, continued work will most likely reveal their broad influence on Fe cycling in marine and saline terrestrial environments.

### Essential Methods

#### *M. ferrooxydans* PV-1 Fe oxidation kinetics

Prior to each kinetics experiment, fresh PV-1 cells were grown up overnight (approx. 19 hours) on Fe(0) plates following Emerson and Floyd (19). For each 20 mL electrochemistry cell, between 1 and 13 mL of cell suspension were added to fresh ASW (avg. 4.1•10^4^ cells mL^-1^). Following this, the electrochemistry cell was set up with O_2_ maintained at the desired concentrations using a gas regulator (air:N_2_ gas mix) with continual stirring. At the start of each 1-2 hour experiment, 300 μL of 10.6 mM ferrous ammonium sulfate was added to the electrochemistry cell (final concentration 154 μM Fe(II)). Fe(II) was measured throughout the experiment using cyclic voltammetry following Luther et al. (107). At the end of each run, abiotic oxidation was measured by killing the cells with 1 mM sodium azide, adding additional ferrous ammonium sulfate, and running the experiment again. At the end of the biotic and abiotic runs, cells were fixed in a 1.6% paraformaldehyde solution before being Syto 13-labeled and counted using a Petroff-Hausser counting chamber.

#### ZOTU-association networks

For the ZOTU network in Figure 7, all full and partial length sequences from the SILVA database (release 128) were included (85). For the ZOTU network in Figure S3, only discrete Fe mat samples (50-75 mL) were included (32, 43, 47, 51, 93, 95. Network connections were assigned using ZetaHunter (2), which produces edge and node files as an output. ZOTU-association network visualization was done in Cytoscape version 3.4.0. Figure 7 was organized spatially using edge-weighted spring embedded layout, which caused ZOTUs found together in multiple samples to be clustered closer together. Figure S3 was organized spatially using attribute circle layout, which placed ZOTUs with a similar dominant sample type close together.

## Acknowledgements

S.M.M. and C.S.C. drafted the manuscript. All authors contributed to editing. A.G., G.W.L., and C.S.C. designed and implemented *M. ferrooxydans* PV-1 kinetics experiments. S.M.M. and R.M.M. developed ZOTU assignments and networks.

We would like to thank Jarrod Scott and Erin Field for their help on structuring and improving this manuscript. This work was funded by NSF OCE-1155290 (C.S.C.), NASA Exobiology NNX12AG20G (D.E., G.W.L, C.S.C) and NNX15AM11G (D.E.), NSF MGG OCE-1558712 (G.W.L.), Office of Naval Research N00014-17-1-2640 (C.S.C) and N00014-17-1-2641 (D.E.), Agriculture and Food Research Initiative grant number 2012-68003-30155 from the USDA National Institute of Food and Agriculture (R.M.M.), and a University of Delaware Doctoral Dissertation Fellowship (S.M.M.). Computational infrastructure support by the University of Delaware CBCB Core Facility funded by Delaware INBRE (grant number NIH P20 GM103446) and the Delaware Biotechnology Institute. The authors declare no conflict of interest.

## References in tables only

(110) – **Makita et al., 2017**

(111) – **Laufer et al., 2017**

(112) – **Makita et al., 2016**

(113) – **Hassenrück et al., 2016**

(114) – **Gonnella et al., 2016**

